# Baclofen, a GABA_B_ receptor agonist, impairs motor learning in healthy people and changes inhibitory dynamics in motor areas

**DOI:** 10.1101/2025.03.27.645701

**Authors:** Ioana-Florentina Grigoras, Elias Geist, Ainslie Johnstone, William T Clarke, Uzay Emir, Caroline Nettekoven, Jacob M. Levenstein, Liliana Capitao, Charlotte J Stagg

## Abstract

Inhibition mediated by γ-aminobutyric acid (GABA) is implicated in motor plasticity and learning, with [GABA] in the motor cortex decreasing during motor learning. However, the causal relationship between [GABA] and learning has yet to be determined. Here, we conducted a within-subject, double-blind, placebo-controlled, crossover study to investigate the effect of increased GABAergic inhibition via GABA_B_-receptor agonist baclofen on motor learning and Magnetic Resonance Spectroscopic Imaging (MRSI) metrics. Increasing GABA-mediated inhibition with baclofen did not change response times, but significantly impaired motor sequence learning. In parallel, we demonstrated a blunting of the expected decrease in [GABA] during motor learning. These results highlight a causal role for GABAergic inhibition in motor learning and may have clinical implications: baclofen is commonly used to treat post brain-injury spasticity, but may impair motor learning during rehabilitation.

## Introduction

Motor learning is essential in everyday life: from learning how to reach and grab objects to learning complex movements, our daily activities rely on previously acquired motor skills. Increasing evidence indicates that inhibition mediated by γ-aminobutyric acid (GABA), the most common inhibitory neurotransmitter, plays a key role in motor plasticity and learning^1,2^. Early mechanistic studies in rodents showed that GABA antagonists in the primary motor cortex (M1) induced long-term potentiation (LTP)-like plasticity^3^, while TMS studies in humans indicate that administration of GABA agonists leads to the suppression of LTP-like processes in M1^4^. Further, the concentration of GABA ([GABA]) in M1 decreases during motor learning^5^ and decreases in M1 [GABA] induced by non-invasive brain stimulation correlate with better motor learning^6^. However, it is not clear whether this observed decrease in GABA is *necessary* for motor learning.

Baclofen is a specific GABA_B_-receptor agonist commonly used in clinical practice as a muscle relaxant^7^. When administered orally, it crosses the blood-brain barrier and reaches a peak plasma concentration in approximately 1-2 hours, with a half-life of 3-6 hours^8^. Its GABA_B_-receptor specificity makes it a good pharmacological intervention for motor learning studies, as GABA_B_ receptors show high expression in areas related to motor control, such as the frontal cortex, thalamic nuclei and cerebellum^9^. Indeed, impairments in visuomotor adaptation after a single dose of 10-20mg baclofen have previously been reported^10,11^. However, the neural mechanisms underlying this behavioural change remain unclear.

Eighteen young healthy participants participated in a within-subject, double-blind, placebo-controlled pharmaco-MRI study. Each participant had an MRI brain scan 45 minutes after administration of a single dose of 20mg of baclofen or placebo, during which Magnetic Resonance Spectroscopic Imaging (MRSI) data was acquired from the primary motor cortex (M1) and premotor cortex (PMC) before and after learning of a serial reaction time task (SRTT). We hypothesised that baclofen would reduce motor learning compared with placebo, and that behavioural decrement would be accompanied by a blunting of the expected learning-related [GABA] decrease in the motor cortices contralateral to the hand performing the task.

## Results

### Baclofen significantly impairs motor sequence learning, but not motor performance

To quantify motor learning, participants performed a serial reaction time task (SRTT) with their right hand whilst in the MRI. Response times (RTs) in the first block (R1), where stimuli were presented in a non-predictable order, were not significantly different between baclofen and placebo sessions (paired t-test: t(13) = 0.161, p = 0.875, Figure 1A), suggesting that baclofen did not modulate motor performance *per se*. We therefore normalised the median RT for each sequence block (S1 to S11) to the median RT in block R1 to directly compare learning between the two sessions.

**Fig. 1.**
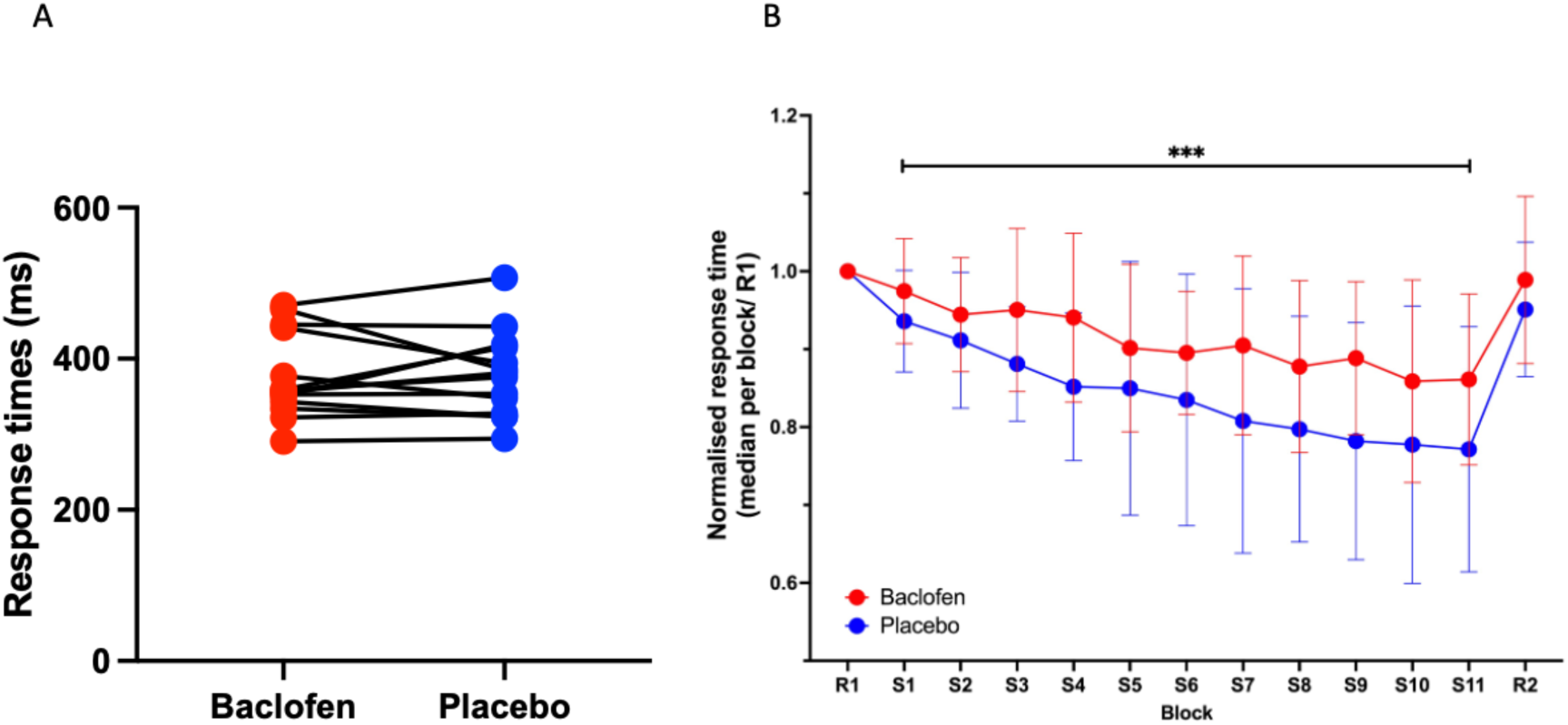
The effect of baclofen on the serial reaction time task (SRTT) (A) No significant differences between baclofen and placebo sessions for response times on the first random block (B) Motor sequence learning was significantly impaired after baclofen administration compared to placebo (R1 and R2 – random blocks 1 and 2; S1 to S11 – sequence blocks 1 to 11)

Consistent with learning, participants significantly reduced their RTs during the task blocks where visual stimuli were presented in a learnable sequence [Repeated-Measures ANOVA with within-subject factors of Drug (baclofen, placebo) and Block (S1 to S11), Main Effect of Block (F(3.179, 41.330) = 9.261, p < 0.001)]. As previously reported, baclofen significantly reduced learning compared with placebo [Main Effect of Drug (F(1,13) = 6.070, p = 0.029), Figure 1B), but there was no significant Block by Drug interaction (Block x Drug Interaction (F(4.568, 59.380) = 1.089, p = 0.373)).

### Baclofen blunts learning-related [GABA] decrease during motor learning

To investigate the neurochemical effects of Baclofen, we quantified mean [GABA] in M1 and PMC before and after participants performed the SRTT (Figure 2A-B). The predicted [GABA] decrease during motor learning was significantly different between the baclofen and placebo sessions, such that baclofen led to a blunting of the expected learning-related decrease in GABA [Repeated-measured ANOVA with factors of Drug (baclofen, placebo), Time (pre-task, post-task) and Region of Interest (ROI; left M1, left PMC), Drug x Time interaction (F(1, 13) = 5.908; p = 0.030; Figure 2C)].

**Fig. 2.**
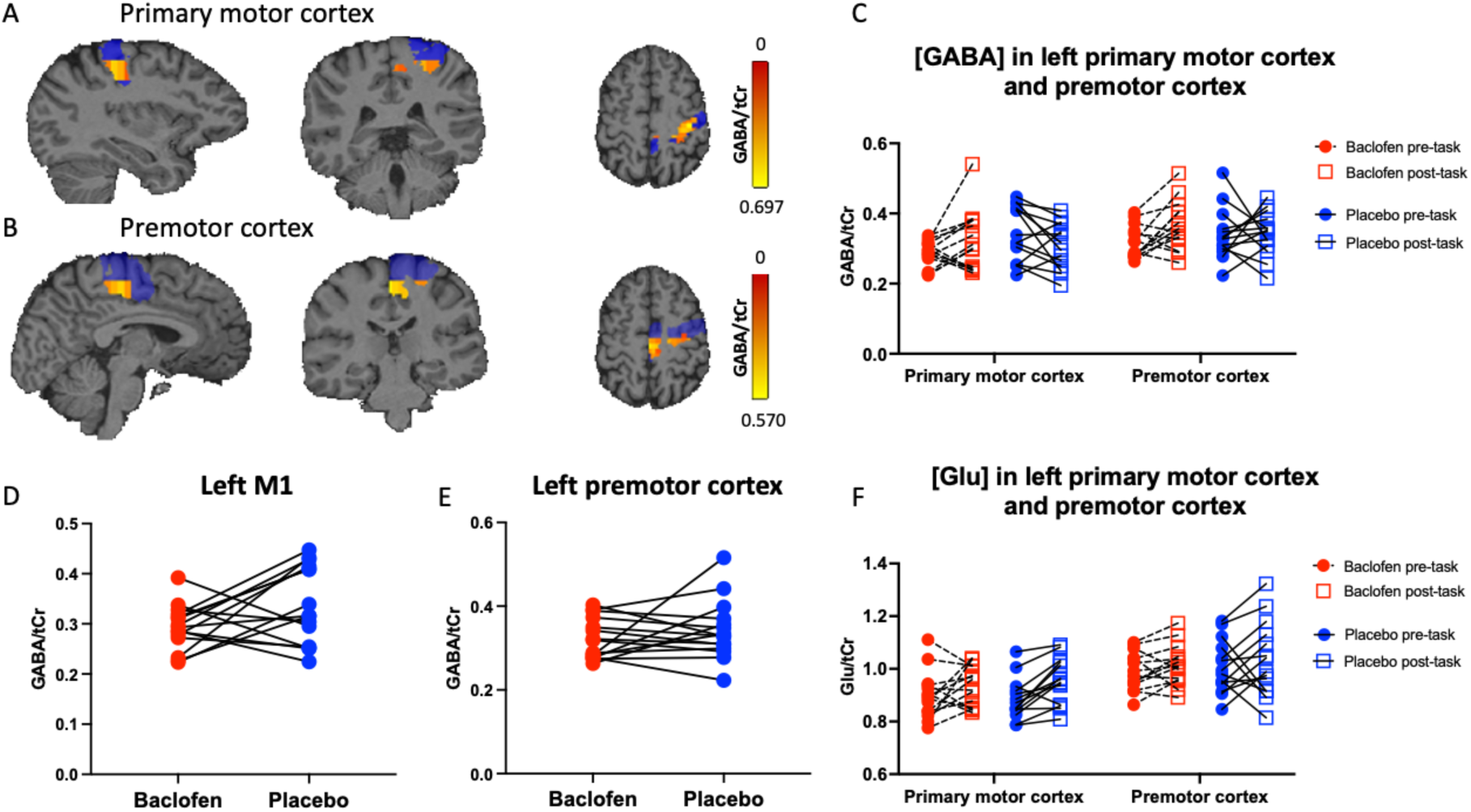
(A-B) Examples of GABA/tCr concentrations in voxels in the left primary motor cortex (A) and left premotor cortex (B). The anatomical mask for each brain region is shown in blue. (C) [GABA] before and after motor learning task. Baclofen significantly affected the pattern of [GABA] change during motor learning across both ROIs (D-E) Pre-task [GABA] did not significantly change in the left primary motor cortex (D) and left premotor cortex (E) after baclofen administration. (F) [Glu] before and after motor learning task. Baclofen did not significantly affect the pattern of [Glu] change during motor learning, but there was a significant increase in [Glu] between the pre- and post-task timepoints.

To determine whether this effect of baclofen on learning-related [GABA] was driven by differences in pre-task [GABA] between baclofen and placebo in left M1 and premotor cortex, we compared [GABA] prior to learning. There was no significant difference in [GABA] between the baclofen and placebo sessions in either region prior to learning (M1: t(14) = 1.915, p = 0.076; PMC: t(14) = 0.946, p = 0.360, Figure 2D-E). ).

To determine the anatomical specificity of this baclofen-induced blunting of learning-related [GABA] decrease, we performed identical analyses for the right, ipsilateral, M1 and PMC (Repeated-measured ANOVA with factors of Drug (baclofen, placebo), Time (pre-task, post-task) and ROI (right M1, right PMC)). Unlike the left, contralateral, cortical areas, there was no significant Drug x Time interaction for the homologous right hemisphere regions (F(1, 13) = 0.026; p = 0.874).

To determine the neurochemical specificity of our [GABA] results, we performed identical analyses for glutamate in the left M1 and PMC (Figure 2F). Unlike for [GABA], there was no significant Drug x Time interaction for glutamate (F(1,13) = 0.579; p = 0.461). There was a significant main effect of Time (F(0.840, 10.920) = 8.174; p = 0.019), suggesting, as might be expected, that glutamate increased in both brain regions after learning, but this increase did not differ between drug conditions.

### MRSI-derived [GABA] measurements were significantly correlated with behavioural metrics

Finally, we investigated whether [GABA] related to behavioural metrics. Mean RT during the random blocks of the SRTT was significantly correlated with pre-task [GABA] in the left PMC in the placebo session, such that greater left PMC [GABA] correlated with slower RTs (r(13) = 0.539, p = 0.038; figure 3A). This relationship was not demonstrated in the baclofen session (r(13) = -0.230, p = 0.409; r-to-z (placebo v baclofen) *z =* 2.050, *p =* 0.040). There were no significant correlations between pre-task M1 [GABA] and RTs in either session (placebo: r(13) = 0.274, p = 0.324; baclofen: r(13) = -0.089, p = 0.752, Figure 3B). This relationship was neurotransmitter-specific: we found no significant correlations between RTs and glutamate levels in either the left M1 (placebo: r(13) = 0.253, p = 0.363; baclofen: r(13) = -0.312, p = 0.258) or the left PMC (placebo: r(13) = 0.019, p = 0.946; baclofen: r(13) = -0.188, p = 0.502).

**Fig. 3.**
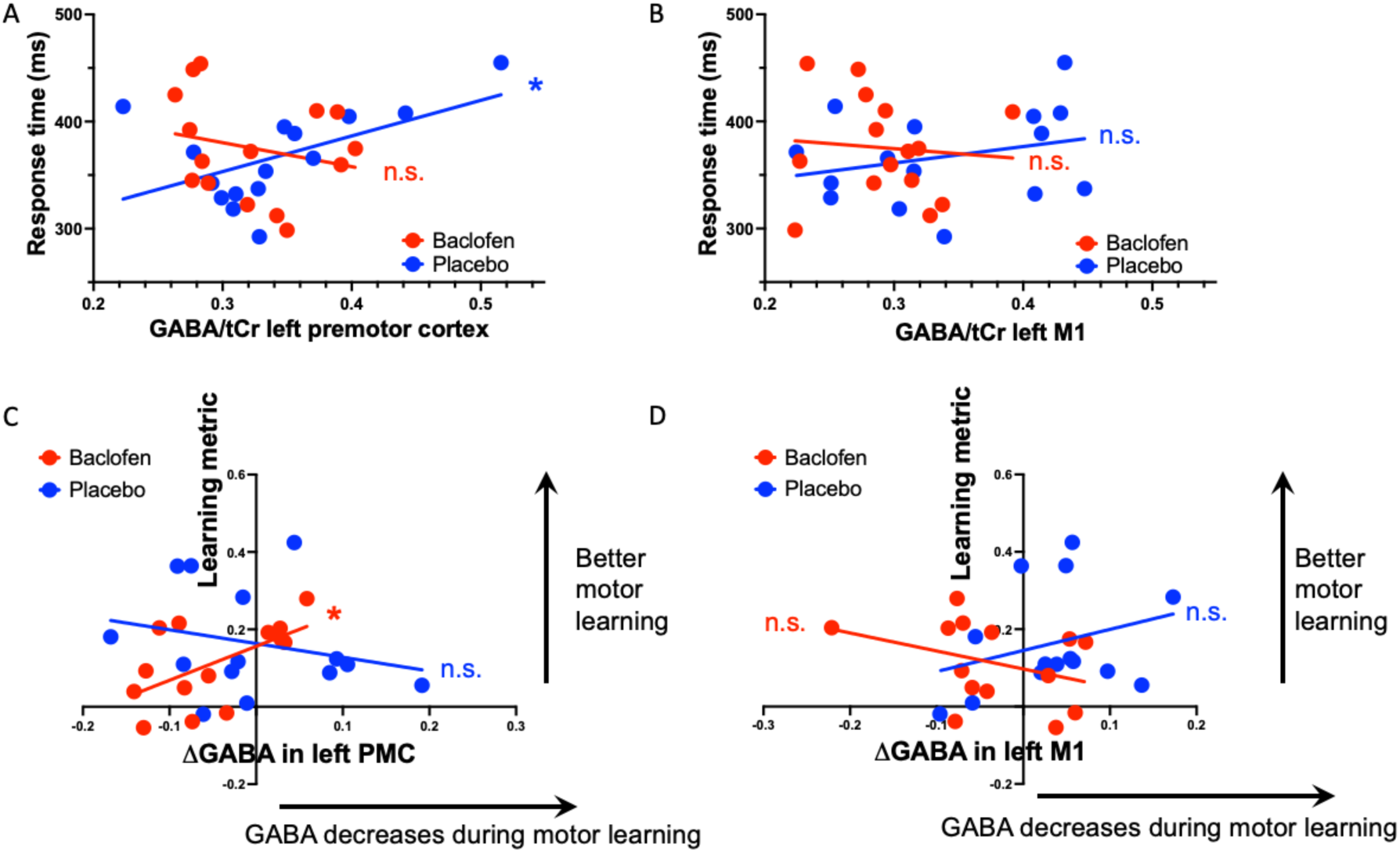
Relationship between pre-task [GABA] and response times during random button presses. (A) Significant correlation between pre-task [GABA] in the left premotor cortex (A) and response times during the placebo session. (B) No significant correlation between pre-task [GABA] in the left M1 and response times (C-D) Relationship between Δ[GABA] and a learning metric calculated based on performance during the SRTT

We then went on to investigate the relationships between neurotransmitters and motor learning. There was a significant correlation between the learning-related change in [GABA] in left PMC and the learning-related decrease in RT in the baclofen session, but not in the placebo session, such that greater decreases in PMC [GABA] correlated with greater learning (baclofen: r(12) = 0.576, p = 0.031; placebo: r(12) = -0.245, p = 0.399; r-to-z (baclofen v placebo) *z =* 2.140, two-tailed *p =* 0.032, Figure 3C). There was no such relationship between learning and change in left M1 [GABA] (baclofen: r(12) = -0.348, p = 0.223; placebo: r(12) = 0.288, p = 0.318, Figure 3D). There was a significant difference between the [GABA] change and behavioural change correlations in the left PMC and M1 on the baclofen sessions (r-to-z *z =* 2.390, two-tailed *p =* 0.017).

When looking at glutamate, we found a significant correlation between learning-related change in [Glu] in the left M1 and the learning-related decrease in RT in the baclofen session, but not in the placebo session. Specifically, smaller increases in M1 [Glu] correlated with greater learning (baclofen: r(12) = 0.571, p = 0.033; placebo: r(12) = -0.076, p = 0.797). There was no such relationship between learning and change in PMC [Glu] (baclofen: r(12) = -0.047, p = 0.873; placebo: r(12) = 0.285, p = 0.323).

### Baclofen increased contentedness, but did not significantly alter working memory

To ensure that the demonstrated baclofen-induced decrease in motor learning was not driven by changes in mood, working memory or attention, participants performed a series of tests.

The 16 Bond-Lader Visual Analogue scales (BLVAS) were loaded onto three mood components: alertness, contentedness, and calmness. Each mood component was quantified before (baseline), and 1h and 2.5 hours after drug administration. There was a significant main effect of drug on contentedness, but not on alertness or calmness (Main Effect of Drug; Contentedness: F(1, 14) = 5.142, p = 0.038; Alertness: F(1, 14) = 4.164, p = 0.061; Calmness: F(1, 14) = 0.653, p = 0.433), such that participants felt more content in the baclofen than placebo session.

The possible effects of baclofen on cognition were assessed via the CANTAB battery. There was no significant drug-related difference in task performance on the spatial working memory task (Main effect of Drug (F(1, 17) = 0.335, p = 0.570)), the pattern recognition memory task (Main effect of Drug (Accuracy: F(1, 17) = 1.294, p = 0.271; RT: F(1, 17) = 1.792, p = 0.198)), Rapid visual processing (paired t-test RT: t(15) = 1.759, p = 0.099) or Spatial Span (Main effect of Drug (Length: F(1, 17) = 2.227, p = 0.154; Errors: (F(1, 17) = 0.221, p = 0.644)).

## Discussion

Understanding the physiological changes underpinning motor learning is essential if we are to develop novel approaches to enhance plasticity, and hence optimise behaviour. Here, we used the GABA_B_ receptor-specific agonist baclofen to address the hypothesis that decreasing GABA is necessary for optimal motor learning. We found that increasing GABA-mediated inhibition via baclofen led to significantly impaired learning, in the absence of changes in response times generally. Baclofen blunted the expected learning-related decrease in [GABA], and the decrease in PMC [GABA] during learning correlated with decreased RTs in the baclofen session, but not during placebo. There were no drug-induced changes in working memory that would explain these drug-related effects.

### Baclofen impairs motor sequence learning

Since [GABA] decreases in M1 are associated with motor learning^5,6,12^ and baclofen is known to suppress neuroplastic processes in the motor cortex^4^, we tested whether increasing GABAergic inhibition using a pharmacological intervention would impair motor learning in healthy volunteers. Baclofen led to a significant decrease in motor learning, supporting the hypothesis that decreases in [GABA] are necessary for motor learning to occur. To the best of our knowledge, this is the first report of impairments in motor sequence learning following baclofen administration: our previous paper^11^, using 10mg of baclofen demonstrated a significant baclofen-induced decrement in visuomotor adaptation, but only a trend towards impaired sequence learning. Differences in drug dose and task design might account for this discrepancy, in particular, our previous paper used 10mg baclofen rather than the 20mg used here.

### Baclofen modulates learning-related changes, but not baseline, [GABA]

Intuitively, we might have expected that resting MRS(I)-derived metrics of GABA might be modulated by baclofen. However, previous studies of GABA-modulating drugs have not shown consistent modulation of MRS-quantified GABA in cortical areas in humans, most reporting null results^13–18^. This is likely explained by MRS metrics not directly measuring the synaptic activity of GABA receptors^19–22^. Instead, MRSI provides measurements of the total amount of MR-visible GABA within each of the 96 MRSI voxels (375 mm^3^), acquired over minutes. This MRSI-derived GABA measurement has been hypothesised to reflect extrasynaptic, tonic inhibition^19^, although this is yet to be confirmed.

Baclofen has non-subtype-specificity for any GABA_B_ receptor subtype, activating both pre- and post-synaptic GABA_B_ receptors^23^. Presynaptically, baclofen binds to the GABA_B_ receptor and decreases the quantity of neurotransmitter released into the synaptic cleft. Postsynaptically, baclofen binds to the GABA_B_ receptor and leads to hyperpolarisation of the cell, but its binding to the receptors instead of the endogenous neurotransmitter could mean that the endogenous GABA remains longer in the synaptic cleft until feedback mechanisms increase its reuptake into glial cells and neurons. Therefore, depending on the proportion of pre-and post-synaptic receptors bound by baclofen, total [GABA] might be expected to either decrease or increase^24–27^. Only one study has so far looked at the effect of baclofen on MRS-derived metrics and reported no significant changes in the right parietal lobe [GABA] between baclofen and placebo^17^. However, this study used a between-subject design in individuals with alcohol-dependence who received a 2-week treatment with baclofen at a high dose (30-75mg), which makes it difficult to predict what the effect of a single-dose of baclofen would be in a healthy population.

### Baclofen-induced changes in PMC [GABA] significantly correlate with changes in motor sequence learning

Baclofen blunts the physiological M1 [GABA] decrease during motor learning, which was associated with impairments in performance on a motor sequence learning task. We found a significant relationship between learning-related changes in the premotor cortex [GABA] and learning-related changes during SRTT during the baclofen session, but not placebo such that, better learning was associated with a larger decrease in the premotor cortex [GABA] during motor learning.

Since the premotor cortex is involved in motor planning and movement preparation, it is not surprising that it has an important contribution towards motor learning^28^, being linked to associative learning (sensory cues become associated with appropriate motor commands) in both healthy individuals and patients^29–31^, as well as implicit motor sequence learning and imitation learning^32,33^. Moreover, reorganisation of the premotor cortex was associated with improvements in motor function after stroke^34^.

Higher task fMRI-derived activation of the dorsal PMC during motor learning has also been associated with higher cognitive demands of the task^35^ and better behavioural outcomes on a motor learning task^36^. Therefore, it is possible that during the baclofen session, when GABAergic inhibition is higher in M1, the premotor cortex contributes more to the process of motor learning than in the placebo session, but that hypothesis remains to be tested.

In the placebo sessions, there were no significant correlations between learning-related [GABA] changes in either M1 or PMC [GABA] and learning-related behavioural changes, which is consistent with previous findings^37^. One possible explanation is that [GABA] dynamics and behavioural improvements do not follow a linear relationship. Alternatively, by collecting the MRSI data in the 10 minutes following the motor sequence learning task, [GABA] levels might be returning to baseline and thus, our [GABA] metrics might be affected by homeostatic processes not related to motor learning.

### Relationship between response times and [GABA] in left PMC in placebo sessions

We found a significant correlation between RTs and [GABA] in the left premotor cortex during the placebo sessions, but not after baclofen administration. The premotor cortex is known to be involved in motor planning and sensorimotor integration and has strong connections to the ipsilateral M1^28,38–41^. fMRI studies investigating the brain areas active during simple response time tasks show a cortical network linked to the identification of the visual stimuli and movement execution, which includes the bilateral premotor cortices^42^.

In non-human primates, micro-stimulation of the premotor cortex, but not motor cortex, during the preparatory activity, led to slower response time in monkeys, likely because the cortical area needs time to restore preparatory activity^43^. In humans, delivering single TMS pulses to the premotor cortex at less than 25ms after the visual stimulus leads to faster RTs, suggesting that preconditioning the premotor cortex decreases the time needed for motor execution^44^. Our results are in line with these studies, showing that lower inhibition in the premotor cortex is associated with faster response times; individual variability in [GABA] may be the neural substrate for individual variability in response times.

Previously, our group has demonstrated a correlation between baseline RTs and M1 [GABA]^19^, which was not demonstrated here. However, previous studies have used a much larger voxel volume (i.e., 8 cm^3^ as opposed to the 0.375 cm^3^ used here). Therefore, although previous studies used an MRS voxel focused on M1, that voxel included regions of the premotor cortex, which might have been driving the correlations previously observed.

### [Glu] in motor areas increases after motor learning

Finally, we found a significant increase in glutamate after the motor learning task across the two study sessions, which likely reflects the increased activity in the motor areas during motor task performance. The possible mechanisms underlying this process have been previously discussed in detail, although definitive evidence confirming these ideas is still lacking^45^. Generally, intermediate to long echo times (>20ms), as used here (TE=32ms), have been proposed as more sensitive to compartmental shift^46,47^ than shorter echo times, so the increase in glutamate after the task could potentially be a result of glutamate moving from the vesicular compartment, largely invisible to MRS, to the extracellular and cytosolic compartment, visible to MRS.

Other studies have also reported increases in glutamate after motor activation (finger tapping, hand clenching) and motor learning, using sequences with TE ranging from 12ms to 80ms^48–50^ and a meta-analysis investigating changes in glutamate after motor and visual task performance is consistent with our findings^46^. Thus, the changes we found here might not be specific to motor learning, but motor activation in general.

We also found a significant relationship between learning-related changes in the left M1 glutamate and learning-related changes in behavioural metrics in the baclofen session, such that greater increases in glutamate are associated with lower learning. It is possible that, with increased inhibition *via* baclofen, the proposed “excitation:inhibition balance” changes and higher increases in glutamate become maladaptive. Some previous studies report no significant correlation between glutamate changes and learning improvements^50,51^, which is consistent with our results in the placebo session. However, there is also a report of resting [Glu] levels predicting learning, although there are no reports of correlations between change in [Glu] and learning improvements^52^.

## Conclusion

We conducted a within-subject, double-blind, placebo-controlled study to investigate how increasing GABAergic inhibition via the pharmacological agent baclofen affected motor learning and brain chemistry. We found that baclofen significantly impaired performance on a motor sequence learning task, but did not impair simple response times. These behavioural impairments were associated with significant learning-related changes in [GABA] in the left M1 and left premotor cortex.

In previous studies in older healthy individuals, motor learning was not associated with decreases in [GABA], but learning-related changes in GABA significantly correlated with motor learning improvements, age, and baseline GABA^53^. Stroke survivors with a previous history of receiving GABA agonists had a significantly worse motor function on admission to a rehabilitation program, though the administration of GABA agonists had no significant effect on subsequent motor rehabilitation outcomes^54^.

Taken together, these results may inform a potential change in clinical guidance towards baclofen use: if baclofen administration impairs motor learning in healthy people, further studies should investigate whether it also impairs motor rehabilitation in patients who receive it as a muscle relaxant.

## Methods

### Participants

We recruited 18 healthy participants who provided their written informed consent to all experimental procedures, as approved by the Central University Research Ethics Committee (Ethics reference: R55534/RE004). All participants met the inclusion criteria: aged 18-35 years (mean age +/- SEM 24.3 +/- 4.1 years, 8 males), right-handed as per the Edinburgh Inventory^55^, no self-reported history of any psychiatric or neurological illness, not taking any medication, not musically trained (not more than Grade 6 on the Associated Board of the Royal Schools of Music) and 3T MRI safety criteria.

### Experimental design

This was a within-subject, double-blind, placebo-controlled study. Participants attended two sessions at least one week apart, starting at the same time of day, with the order of the sessions randomised across the group (Figure 4A).

**Fig. 4.**
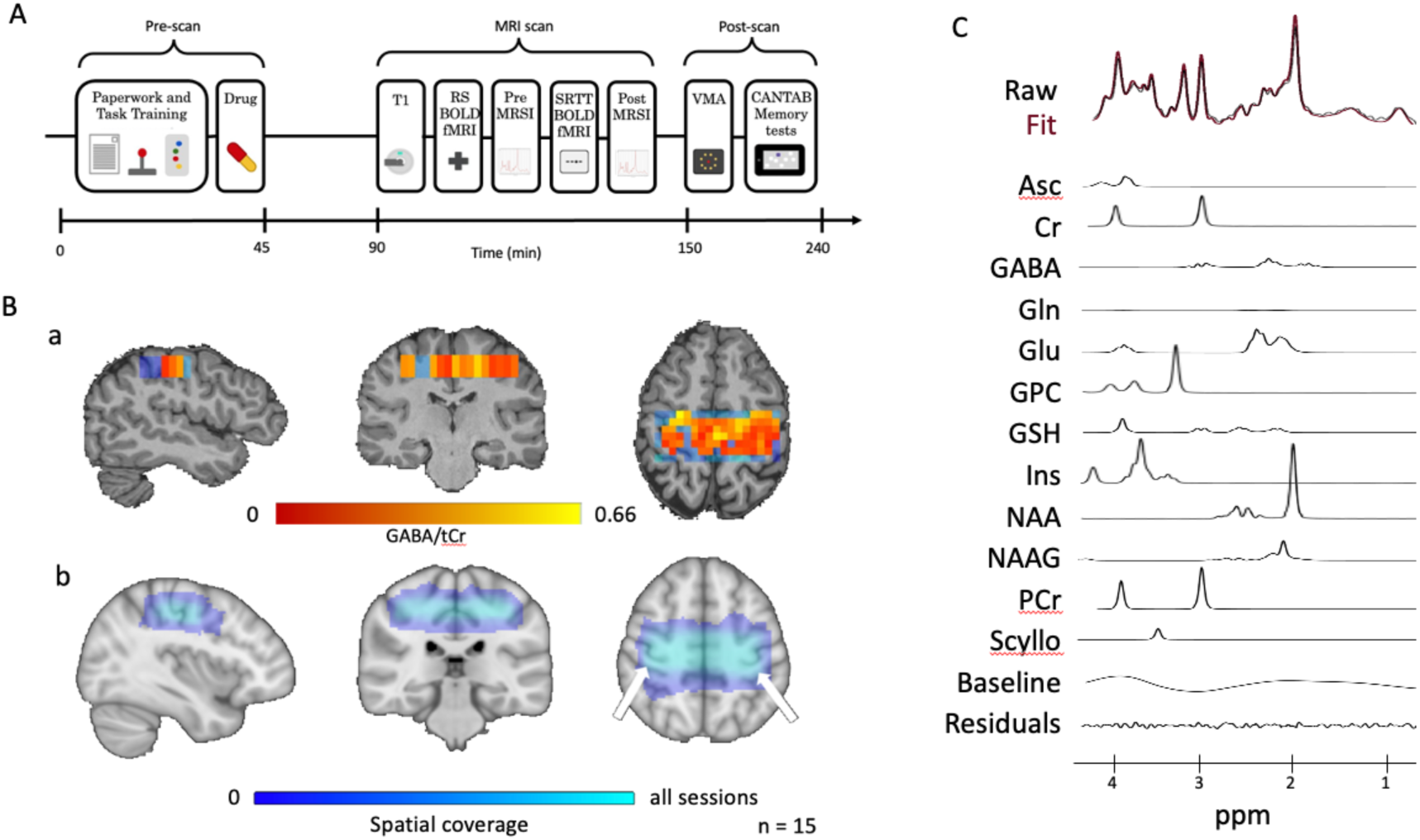
(A) Timeline of each testing session. (B) a. Map of GABA/tCr in one of the sessions. Voxels that pass quality checks are shown in red and excluded ones in blue. b. Group level map of the spatial coverage of the MRSI volume on interest across all sessions (15 participants x 2 session); (C) Representative spectrum

On arrival, participants completed a Bond-Lader Visual Analogue Scale (BLVAS) mood questionnaire and were familiarised with the serial reaction time task (SRTT). Participants were then given the drug (baclofen or placebo) and had a 45-minute break. Participants then filled in a second BLVAS questionnaire before having an MRI scan. We first acquired a T1- weighted structural image before pre-task MRSI (sequence details below), which started approximately 75 minutes after drug administration, to allow for baclofen to be absorbed and reach peak plasma concentration^8,56^. Participants then completed the SRTT, followed by a post-task MRSI acquisition. The scan timings were chosen so that the MRSI scans and motor learning tasks fell within the peak plasma concentration time for baclofen, which is 1-2 hours after drug administration^8^. After the MRI scan, participants filled in a third BLVAS questionnaire 2 hours after treatment administration and performed a battery of Cambridge Cognition (CANTAB) memory tests.

### Serial Reaction Time Task (SRTT)

Participants were instructed to use their right index, middle, ring, and little fingers to press buttons 1-4 on an MRI-compatible 4-button box in response to a visual cue, as quickly and accurately as possible. Participants were also instructed to not press a button before the visual cue appeared on the screen. At the beginning of each block, participants would see a screen with four dashes. One of the dashes would then be replaced by an asterisk, which represented the visual stimulus for the participant to press the corresponding button. The SRTT was composed of two types of blocks: sequence blocks and random blocks, each with 60 trials repeated at a frequency of 1.5Hz. During random blocks, the visual cues were presented in a random order. During each sequence block, the same 12-button sequence was repeated five times. A different sequence was used for each session (sequence 1: 3-1-2-4-2-3-2-1-4-1-3-4; sequence 2: 3-4-2-1-3-1-2-4-1-4-3-2).

We calculated response times (RTs) for each block as the time between stimulus presentation and the button press. We excluded incorrect button presses, pre-emptive button presses (RT < 50ms) and outliers (± 2.7 SD from the mean of each block). We then calculated the median RT for each block and normalised it by dividing it by the median RT of the first random block (R1). To correlate behaviour with MRS measures of [GABA], we also calculated the slope of RT change as a learning measure for each session by fitting a linear regression model to the data in sequence blocks 1 to 11. One participant was excluded from further analysis because their learning measure on one session being an outlier on the Grubbs test.

To investigate how changes in neurochemicals relate to changes in behaviour on the motor sequence task, we first calculated the change of a neurochemical during motor learning as the difference between the pre- and post-task [GABA] and [Glu] respectively, so higher values of this metric would represent larger decreases in that neurochemical during motor learning. To assess changes in motor learning, we calculated the motor learning metric as the difference in median response times between the first and last sequence blocks normalised by the median response time to the first random block [(S1-S11)/R1], so that higher values of this metric would represent larger decreases in response time during motor sequence learning and therefore, better learning.

### MR Data Acquisition and Analysis

Participants had an MRI brain scan in a 3T MAGNETOM Prisma system (Siemens Healthineers, Elangen, Germany) equipped with a 32-channel receive head coil (Siemens Healthineers). Three participants did not complete the MRI scan and were therefore not included in the neuroimaging analysis.

First, we acquired a T1-weighted structural image (MPRAGE, 1 mm isotropic, TR = 1.9 s, TE = 3.96 ms, TI = 912 ms, TA = 7.3 min, field of view 232 x 256 x 192mm^3^, flip angle 8°). MRSI was acquired pre- and post-task using a semi-LASER prepared MRSI sequence, with density-weighted concentric ring trajectories (CRTs) k-space sampling^57^, TR = 1.4 s, TE = 32 ms, TA = 2 x 4.5min, voxel size = 5 x 5 x 15 mm^3^, field of view = 85 x 35 mm^2^ , slice thickness = 15 mm). The semi-LASER selected volume (“MRSI slab”) was manually placed to cover both the left- and right-hand knobs at the posterior margin of the precentral gyrus^58^, excluding tissue outside the brain to minimise contamination due to mobile lipids ^57^.

Reconstruction of MRSI data and neurochemical quantification was performed using in-house scripts and LCModel, as described in previous studies^57,59–61^. Briefly, after metabolite cycling reconstruction^62^ and coil-combination^63^, we corrected for frequency and phase shifts, removed residual water using HLSVD^64^, and corrected for eddy currents^65^. Neurochemicals were quantified using LCModel with a basis set containing 26 metabolites, default LCModel macromolecules, no soft constraints on metabolites, a baseline stiffness setting (DKMTMN) of 0.25 and a chemical shift of 0.5 to 4.2ppm. Metabolite concentrations are reported as a ratio to total creatine (creatine + phosphocreatine; tCr). We then excluded voxels with Cramer-Rao lower bands (CRLB) > 50%, signal-to-noise ratio (SNR) < 40 or GABA/tCr > 1 (Fig.4B-a).

To confirm the accurate placement of the MRSI slab in all participants, metabolite maps were registered to MNI space using linear and then non-linear registration (FLINT and FNIRT, ^66,67^). All MRSI slabs included the left-hand knob (Fig.4B-b). A representative spectrum is included in Figure 4C. To quantify GABA in the brain regions responsible for motor control, we registered anatomical MNI maps of left and right M1 and premotor cortex to each participant’s structural scan using FLIRT and FNIRT^66^. For each participant, we calculated the mean [GABA] from the voxels that passed quality control in each region of interest.

### Working memory tests

To investigate the effect of baclofen on cognitive functions, participants completed four working memory tests from the Cambridge Neuropsychological Test Automated Battery (CANTAB) on a tablet (Apple iPad Air 2) at approximately 3 hours after treatment administration^68^. These four tests were: Spatial Working Memory (SWM), Pattern Recognition Memory (PRM), Spatial Span (SSP), and Rapid Visual Processing (RVP). Due to technical difficulties, 3 participants could not complete the SWM, PRM and RVP tests on one of the sessions.

#### Spatial working memory

A number of boxes appeared on the screen and participants were instructed to tap one box to find out whether there was a token in it. Only one token appeared in each of the boxes, so participants had to remember which boxes they had already found tokens in and which boxes they still needed to check. Participants committed errors whenever they tapped a box in which they had already looked for a second time. The number of total errors was reported for increasing task difficulties (4, 6, 8 and 12 blocks). A strategy metric was also recorded, quantifying how many times the participants had started searching from the same block, indicating a strategy for task performance.

#### Pattern Recognition Memory

Participants were presented with a set of shapes, which they were instructed to remember. Pattern recognition memory was evaluated at two time points: immediately after the presentation of the set of shapes and at approximately 20 minutes. Participants were asked to recall the shapes they remembered from two possible options. The percentage of correct choices and the response times for pattern recognition were recorded for both the immediate and delayed time points.

#### Rapid Visual Processing

Digits between 2 and 9 were briefly displayed in the centre of the screen in a pseudorandom order. Participants were instructed to tap a red button on the screen if the following target sequences appeared: 3-5-7, 2-4-6 and 4-6-8. We recorded three metrics: the accuracy of sequence detection, median response times and the probability of false alarm.

#### Spatial Span

For each trial, a set of 9 boxes appeared on the screen, out of which a subset would change colour one at a time, forming a sequence. In the forward version of this task, participants were asked to tap the boxes in the order they had lit up previously; in the reverse version, participants were asked to tap the boxes in the reserve order. The sequence gradually increased from 2 to 9 boxes, making the task more difficult to perform. The length of the most difficult correctly-recalled sequence was recorded, as well as the number of errors made, for both the forward and the reverse spatial span.

#### Mood questionnaires

Participants completed the Bond-Lader Visual Analogue scale (BLVAS) three times during each session. The BLVAS has 16 scales, each 100mm long with the 2 opposite sides of the spectrum at each end, e.g. alert – drowsy, calm – excited^69^. At each time point, participants were asked to mark how they felt at that specific moment on each scale. Similar to previous studies, the 16 scales were grouped into 3 categories: alertness (alert, strong, clear-headed, well-coordinated, energetic, quick-witted, attentive, proficient, interested), contentedness (contented, tranquil, happy, social, friendly) and calmness (calm, relaxed) ^69^. The mean value for each category was calculated at each time point, with lower values indicating stronger feelings on that respective scale.

### Statistical Analyses

All behavioural data were analysed using MATLAB (v. R2020b, MathWorks) and Prism (v.9.3.0, GraphPad Software). MRSI data were analysed using in-house Bash scripts, R (RCoreTeam 2013) and Prism. Analysis was performed using repeated-measures ANOVAs with Geisser-Greenhouse’s correction or paired t-tests with significance levels of 𝛼 = 0.05 to investigate the within-subject effects of baclofen compared to placebo. Relationships between MRSI and behavioural metrics were investigated by calculating Pearson’s correlation coefficients. All graphs show mean ± SD or individual data points unless otherwise indicated.

## Acknowledgements

CJS was supported by a Wellcome Trust Senior Research Fellowship (224430/Z/21/Z). WTC was supported by a Wellcome Trust Career Development Award (225924/Z/22/Z). The Wellcome Centre for Integrative Neuroimaging is supported by core funding from the Wellcome Trust (203139/Z/16/Z and 203139/A/16/Z). WTC was supported by a Wellcome Trust Career Development Award (225924/Z/22/Z). LC currently works at the Psychology Research Centre (PSI/01662), School of Psychology, University of Minho, supported by the Foundation for Science and Technology (FCT) through the Portuguese State Budget (Ref.: UID/PSI/ 01662/2020). LC is individually funded by a Research Fellowship also from the Foundation for Science and Technology (FCT) (Ref: 2021.00415.CEECIND). This study was supported by the NIHR Oxford Health Biomedical Research Centre (OH BRC, NIHR203316). The views expressed are those of the authors and not necessarily those of the NHS, NIHR or the Department of Health.

## Authorship contributions

Conceptualisation: IFG, AJ, CJS

Experimental design and data collection: IFG, EG, AJ, WTC, UE, LC, CJS

Data analysis: IFG, EG, WTC, UE, CN, JML, CJS

Writing – original draft: IFG and CJS

Writing – review & editing: IFG, EG, AJ, WTC, UE, CN, JML, LC, CJS

Funding acquisition: CJS

## Notes

### Competing Interest Statement

The authors have declared no competing interest.

